# Function of the Arabidopsis kinesin-4, FRA1, requires abundant processive motility

**DOI:** 10.1101/107870

**Authors:** Anindya Ganguly, Logan DeMott, Ram Dixit

## Abstract

Processivity is important for kinesins that mediate intracellular transport. Structure-function analyses of N-terminal kinesins have identified several non-motor regions that affect processivity *in vitro*. However, whether these structural elements affect kinesin processivity and function in vivo is not known. Here, we used an Arabidopsis kinesin-4, called Fragile Fiber1 (FRA1), which is thought to mediate vesicle transport to test whether mutations that alter processivity *in vitro* behave similarly in vivo and whether processivity is important for FRA’s function. We generated several FRA1 mutants that differed in their run lengths *in vitro* and then transformed them into the *fra1-5* mutant for complementation and in vivo motility analyses. Our data show that the behavior of processivity mutants in vivo can differ dramatically from *in vitro* properties, underscoring the need to extend structure-function analyses of kinesins in vivo. In addition, we found that high density of processive motility is necessary for FRA1’s physiological function.

**Summary:** This study shows that the motility of kinesin mutants can differ significantly between *in vitro* and *in vivo* conditions and that abundant processive motility is important for FRA1 kinesin’s function.

## INTRODUCTION

Kinesins execute various cellular functions either by powering directional transport of cargo along microtubules or by regulating the dynamics of microtubule ends. Transport kinesins, such as kinesin-1, 2 and 3 members, mediate long-distance transport of diverse cargo including vesicles, protein complexes and nucleic acids (Hall and Hedgecock, 1991; Kanai et al., 2000; Kondo et al., 1994; Okada et al., 1995; Vale et al., 1985; Verhey et al., 2001). While some microtubule regulatory kinesins, such as kinesin-13 members, are non-motile, others such as kinesin-4 and kinesin-8 members are motile but work primarily to regulate the dynamics of microtubule plus-ends (Bringmann et al., 2004; Desai et al., 1999; Gupta et al., 2006; Kurasawa et al., 2004; Varga et al., 2006; Walczak et al., 1996).

Plants contain a large number of kinesins but little is known about which ones are used for transport and microtubule regulatory functions (Richardson et al., 2006; Zhu and Dixit, 2012). Since kinesin-1, 2 and 3 members are absent in plants, other classes of kinesins probably perform plus-end-directed transport functions. Recently, a plant kinesin-4 member, called Fragile Fiber 1 (FRA1), was shown to mediate trafficking of cell wall material along cortical microtubules (Kong et al., 2015; Zhu et al., 2015). Loss of the FRA1 kinesin in a *fra1-5* mutant leads to thinner cell walls, reduced rate of pectin secretion and accumulation of Golgi-associated vesicles, which together indicate that FRA1 likely transports vesicles for secretion. In contrast, both microtubule plus-end dynamics and array organization are unaltered in this mutant (Zhu et al., 2015), demonstrating that unlike metazoan kinesin-4s FRA1 does not regulate microtubule plus-end growth.

Transport kinesins have a common structural organization consisting of a catalytic head domain, a flexible neck linker that mediates inter-head communication and amplifies conformational changes needed for motion, a coiled-coil stalk and cargo binding tail domain. Another shared feature of transport kinesins is their ability to move processively― i.e., take many steps upon binding to a microtubule―which enables efficient long-distance transport by single motors (Howard et al., 1989; Soppina et al., 2014; Yamazaki et al., 1995). Consistent with a transport function, dimeric FRA1 motors have been found to move processively both *in vitro* and *in vivo* (Kong et al., 2015; Zhu and Dixit, 2011; Zhu et al., 2015). To directly investigate the importance of FRA1 processivity for its physiological function, we generated several mutants by modifying different non-motor domains predicted to alter run length and studied them using *in vitro* motility assays as well as genetic complementation and live-imaging in *Arabidopsis thaliana* plants. Our data show that the *in vitro* and *in vivo* motile properties of FRA1 can vary depending on the configuration of the mutant. Further, we found that robust processive motility is critical for FRA1 function.

## RESULTS AND DISCUSSION

### Mutating neck-linker and neck coiled-coil regions alters FRA1 processivity *in vitro*

*In vitro* experiments have shown that processivity of kinesins can be tuned by altering the length and charge of the neck linker domain and the charge of the neck coiled-coil domain (Shastry and Hancock, 2010; Shastry and Hancock, 2011; Thorn et al., 2000). To identify these domains in the FRA1 kinesin, we aligned the amino acid sequences of the motor domains of FRA1 and kinesin-1 to define the start of the neck linker and used the PCOILs and Multicoil programs to predict the start of the neck coiled-coil. These analyses indicated that the neck linker of FRA1 is 14 amino acids, similar to that of human kinesin-1 and kinesin-4 (Fig. 1A).

**Figure 1.**
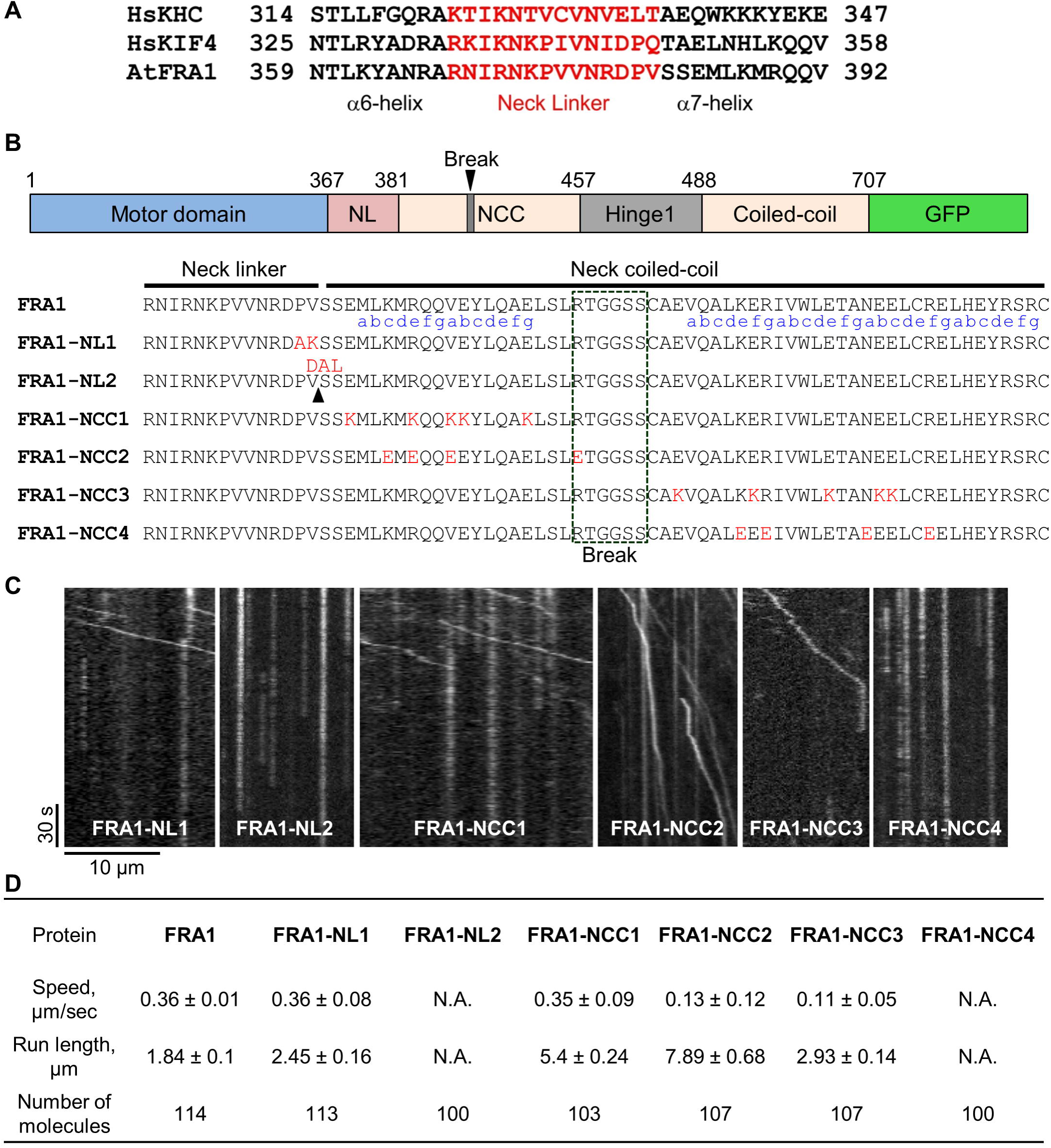

*In vitro* motility analysis of FRA1 neck-linker and neck coiled-coil mutants.

(**A**) Sequence alignment of neck linker and flanking *α*-helices of human kinesin-1 heavy chain (HsKHC), a human kinesin-4 member (HsKIF4) and *Arabidopsis thaliana* FRA1 (AtFRA1). The neck linker regions are shown in red.

(**B**) Schematic representation of the structural domains of the first 707 amino acids of the FRA1 kinesin fused with GFP. The amino acid sequences below show the mutations used in this study in red. The predicted heptad repeats in the neck coiled-coil are shown in blue. NL, neck linker; NCC, neck coiled-coil.

(**C**) Kymographs of the FRA1 mutants from *in vitro* single-molecule motility assays.

(**D**) Motility parameters of wild-type and mutant FRA1 kinesins calculated from the *in vitro* experiments. N.A., not applicable because of lack of directional motility. Values are mean ± SD.

To test whether mutations in the neck linker and neck coiled-coil domains affect FRA1 processivity, we conducted single-molecule *in vitro* motility assays with a Green Fluorescent Protein (GFP)-labeled truncated version of FRA1, called FRA1(707)-GFP, which was previously shown to move processively as a dimer *in vitro* (Zhu and Dixit, 2011). We generated two mutants that changed the length and charge of the neck linker domain: FRA1-NL1, in which the PV residues of the neck linker were substituted to AK, and FRA1-NL2 in which a three amino acid DAL peptide was inserted after the PV residues (Fig. 1B). The NL1 mutation was expected to enhance processivity by creating a straighter, more positively charged neck linker; while the NL2 mutation was expected to reduce processivity by overextending the neck linker (Shastry and Hancock, 2010). We found that FRA1-NL1 had ~ 30% increased processivity and unchanged speed compared to wild-type FRA1, while FRA1-NL2 did not show any processive runs within the spatial resolution limit (~ 250 nm) of our imaging system (Fig. 1C,D).

To test whether the charge of FRA1’s neck coiled-coil domain affects its processivity, we generated four mutants that targeted charged amino-acids either within heptads 1 and 2 or within heptads 3 and 4 (Fig. 1B). Two of these mutants, FRA1-NCC1 and FRA1-NCC3, made the heptads more positively charged by replacing glutamic acid residues with lysine. Conversely, the FRA1-NCC2 and FRA1-NCC4 mutants made the heptads more negatively charged by replacing positively charged amino acids with glutamic acid. Similar to previous findings with human kinesin-1 (Thorn et al., 2000), adding positive charges to the neck coiled-coil domain increased FRA1’s processivity. FRA1-NCC1 showed ~ 3-fold greater processivity and moved at essentially the same speed as wild-type FRA1 (Fig. 1C,D). In contrast, FRA1-NCC3 showed ~ 60% greater processivity and ~ 3-fold lower speed compared to wild-type FRA1 (Fig. 1C,D). One possible reason for why these mutations differentially affect FRA1’s processivity and speed is that they might destabilize the neck coiled-coil to different extents thus affecting the coordination between the two motor domains to varying degree.

Surprisingly, the FRA1-NCC2 mutant showed ~ 4-fold greater processivity even though its neck coiled-coil is more negatively charged than wild-type FRA1 (Fig. 1C,D). This mutant also moved ~ 3-fold slower than wild-type FRA1. Why this mutant has greater processivity than FRA1-NCC1 is not clear and requires further study. In contrast to FRA1-NCC2, the FRA1-NCC4 mutant did not show processive motility under our *in vitro* conditions (Fig. 1C,D). Taken together, these results show that changing the length and charge of the neck linker and neck coiled-coil domains modulates FRA1’s processivity *in vitro* similar to other N-terminal kinesins.

### Functional analysis of FRA1 processivity mutants *in vivo*

To determine if the above mutations affect FRA1 processivity and function *in vivo*, we introduced them in full-length FRA1 labeled with tdTomato and stably transformed these constructs driven by the native *FRA1* promoter into the *fra1-5* knockout mutant.

Expression of wild-type FRA1-tdTomato completely rescued the dwarf phenotype of *fra1-5* plants (Fig. 2A,B). The two neck-linker mutants, FRA1-NL1 and FRA1-NL2, nearly completely rescued the *fra1-5* phenotype, indicating that they are functional (Fig. 2A,B). In contrast, none of the four FRA1-NCC mutants rescued the dwarf phenotype of *fra1-5* in more than 10 independent transgenic lines for each (Fig. 2A,B). This inability was not due to lack of FRA1-NCC transgene expression, as evidenced by real-time RT-PCR measurements of transcript abundance (Fig. 2C).

**Figure 2.**
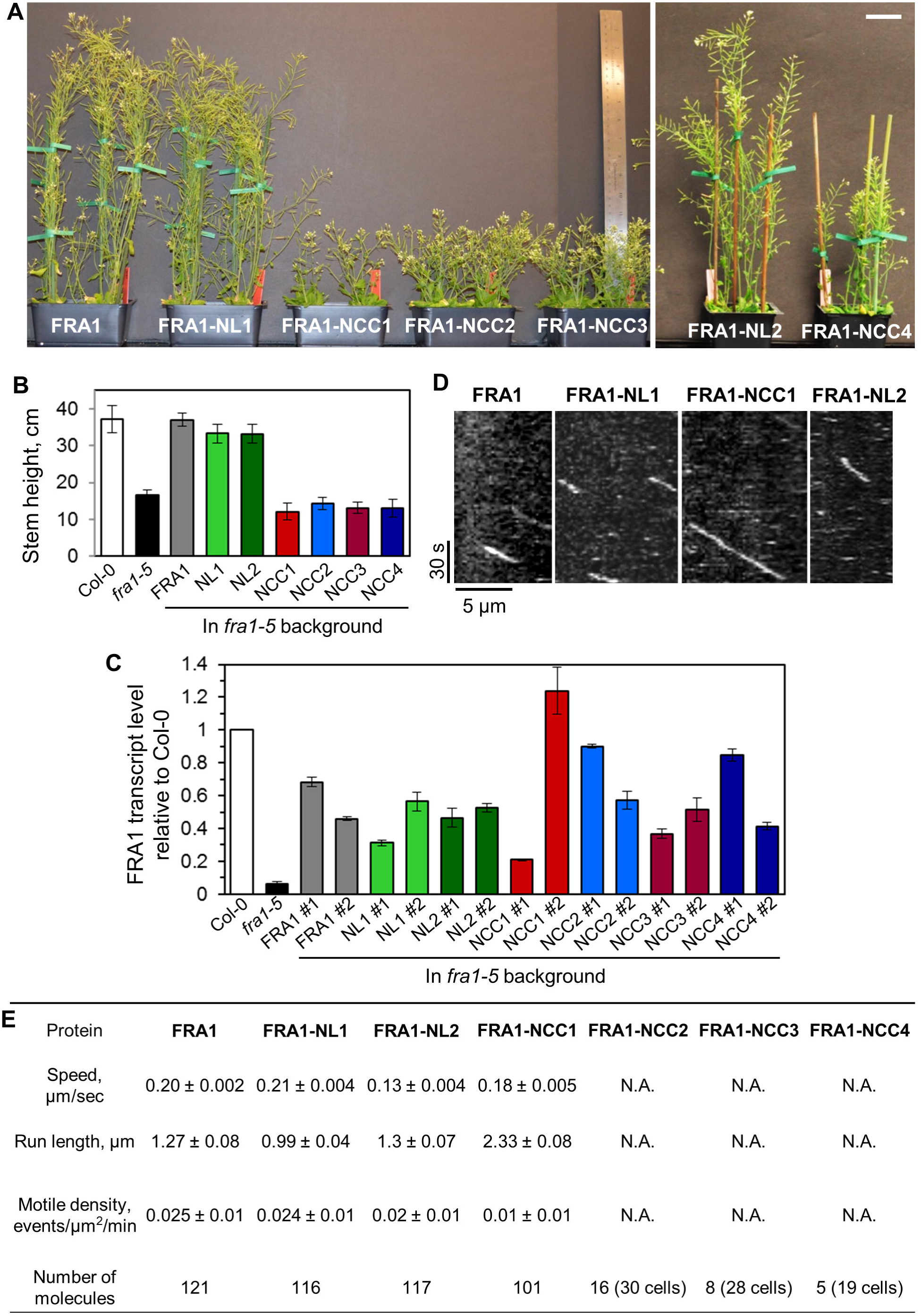

*In vivo* functional and motility analysis of FRA1 neck-linker and neck coiled-coil mutants.

(**A**) Images of 4.5-week old plants expressing FRA1, FRA1-NL1, FRA1-NCC1, FRA1-NCC2, FRA1-NCC3, FRA1-NL2 or FRA1-NCC4 in the *fra1-5* mutant. Scale bar = 5 cm.

(**B**) Heights of primary inflorescence stems. Values are mean ± SD from at least 16 plants for each genotype.

(**C**) *FRA1* expression level relative to wild-type control. Values are mean ± SEM from three biological replicates.

(**D**) Kymographs showing the movement of wild-type FRA1, FRA1-NL1, FRA1-NCC1 and FRA1-NL2 puncta at the cell cortex.

(**E**) Motility parameters of wild-type and mutant FRA1 kinesins determined from live imaging experiments. N.A., not applicable because of extremely rare events of directional motility. Values are mean ± SD.

To determine if *in vivo* functionality of the FRA1 mutants correlates with their motility, we imaged 4-day old hypocotyl epidermal cells using variable angle epifluorescence microscopy. Wild type FRA1-tdTomato puncta moved an average run length of 1.3 ± 0.08 μm at an average speed of 0.2 ± 0.002 μm sec^−1^ (Fig. 2D,E). In contrast to *in vitro* results, the run length of FRA1-NL1 puncta was ~ 25% lower than wild-type FRA1 at 0.99 ± 0.04 μm while their average speed was essentially the same as wild-type FRA1. Furthermore, FRA1-NL2 was highly motile *in vivo* even though it was non-motile *in vitro*. Directionally moving FRA1-NL2 puncta had a run length similar to wild-type FRA1 at 1.3 ± 0.1 µm and an average speed of 0.13 ± 0.01 µm sec^−1^ (Fig. 2D,E). FRA1-NCC1 puncta had ~55% greater run length than wild-type FRA1 at 2.33 ± 0.08 μm and an average speed of 0.18 ± 0.005 μm sec^−1^. Interestingly, even though FRA1-NCC1 is highly processive, it does not complement the *fra1-5* mutant. One potential explanation for this finding is that FRA1-NCC1’s motile density is less than half that of wild-type FRA1 and FRA1-NL1 (Fig. 2E), which would predictably lead to less cargo transport.

The FRA1-NCC2, FRA1-NCC3 and FRA1-NCC4 mutants showed very little directional motility *in vivo* (Fig. 2E) even though FRA1-NCC2 and FRA1-NCC3 were highly processive *in vitro*. For these mutants, we frequently observed static puncta over the ~ 2 min imaging period, indicating that these mutants likely bind to microtubules *in vivo* but lack processive motility. Based on these data, we conclude that robust processive motility is necessary for FRA1’s function in plants.

### Mutation of a conserved serine residue in the FRA1 motor domain alters its motility

Phosphorylation of a conserved serine residue in the motor domain of a *Drosophila melanogaster* kinesin-13, KLP10A, has been shown to decrease its microtubule depolymerizing activity without altering its microtubule binding (Mennella et al., 2009). Sequence alignment with the motor domains of plant FRA1 homologs showed that this serine residue is conserved within the microtubule binding *α*-helix 5 of these kinesins (Fig. 3A). To determine whether this serine potentially plays a role in regulating FRA1 motility, we generated phosphomimetic (FRA1-S334E) and nonphosphorylatable (FRA1-S334A) versions of both FRA1(707)-GFP and full-length FRA1-tdTomato.

**Figure 3.**
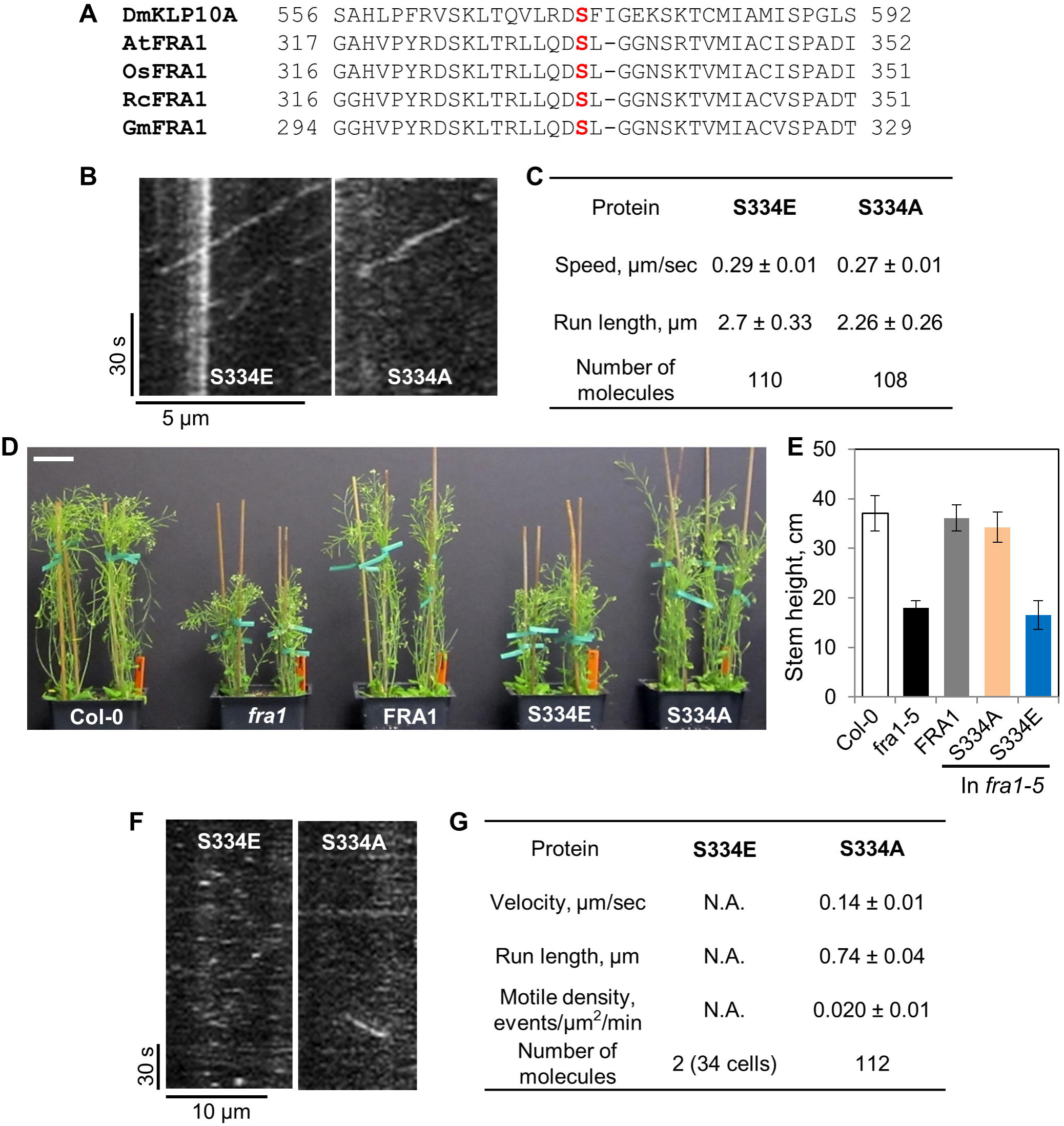

Analysis of FRA1 motor domain phosphorylation mutants.

(**A**) Sequence alignment of the *α*-helix 5 region of the motor domains of Drosophila KLP10A (DmKLP10A) and various plant FRA1 homologs. The conserved serine residue is shown in red. At, *Arabidopsis thaliana*; Os, *Oryza sativa*; Rc, *Ricinus communis*; Gm, *Glycine max*.

(**B**) Kymographs of the FRA1 phosphomutants from *in vitro* single-molecule motility assays.

(**C**) Motility parameters of FRA1 phosphomutants calculated from the *in vitro* experiments. Values are mean ± SD.

(**D**) Images of 4.5-week old plants. Scale bar = 5 cm.

(**E**) Heights of primary inflorescence stems. Values are mean ± SD from at least 16 plants for each genotype.

(**F**) Kymographs showing the movement of FRA1-S334A and FRA1-S334E puncta at the cell cortex.

(**G**) Motility parameters of FRA1-S334A and FRA1-S334E determined from live imaging experiments. N.A., not applicable because of extremely rare events of directional motility. Values are mean ± SD.

*In vitro*, FRA1-S334E and FRA1-S334A showed ~33% and ~22% increased processivity and ~18% and ~23% decreased speed compared to wild type FRA1, respectively (Fig. 3B,C). However, we found that FRA1-S334A was able to rescue the dwarf *fra1-5* phenotype, whereas FRA1-S334E was not (Fig. 3D,E). Live imaging showed that genetic complementation correlated with the motility of these mutant kinesins. Specifically, the FRA1-S334A mutant was motile whereas the FRA1-S334E mutant was non-motile (Fig. 3F,G). Notably, FRA1-S334A is functional even though its average run length and speed *in vivo* are ~ 40% and ~25% less than wild type FRA1 respectively (Fig. 3G). However, the motile density of FRA1-S334A is similar to that of wild type FRA1 (Fig. 3G), indicating that lower kinesin processivity and speed are tolerated as long as sufficient numbers of them are engaged in transport.

### Overexpression of FRA1 phenocopies the *fra1-5* mutant due to gene silencing

Overexpression of FRA1 has been found to phenocopy *fra1* loss-of-function mutants (Zhou et al., 2007), however, the underlying mechanism remains unknown. An increase in kinesin density can decrease processivity *in vitro* by overcrowding the microtubule lattice (Leduc et al., 2012; Telley et al., 2009). Therefore, one possible mechanism for the FRA1 overexpression phenotype is that it creates traffic jams along cortical microtubules, thus reducing processivity and compromising FRA1 function. To test this hypothesis, we expressed wild-type FRA1-tdTomato driven by a constitutive 35S promoter in wild-type *Arabidopsis thaliana* plants. Similar to previous work, our FRA1-overexpression lines mimicked the dwarf stem phenotype of *fra1-5* plants (Fig. 4A,B). However, transcript analysis using real-time RT-PCR revealed that *FRA1* expression was greatly reduced in these plants (Fig. 4C). In addition, immunoblot analysis showed that endogenous FRA1 protein was undetectable in these plants (Fig. 4D). Using live imaging, we also found that FRA1-tdTomato was essentially undetectable at the cell cortex in these plants. Therefore, both endogenous and transgenic *FRA1* expression were suppressed, probably due to 35S-promoter triggered gene silencing (Daxinger et al., 2008; Mishiba et al., 2005; Mlotshwa et al., 2010). These findings explain why FRA1 “overexpression” lines show remarkably similar developmental and cell wall phenotypes as the knockout *fra1-5* line (Zhou et al., 2007).

**Figure 4.**
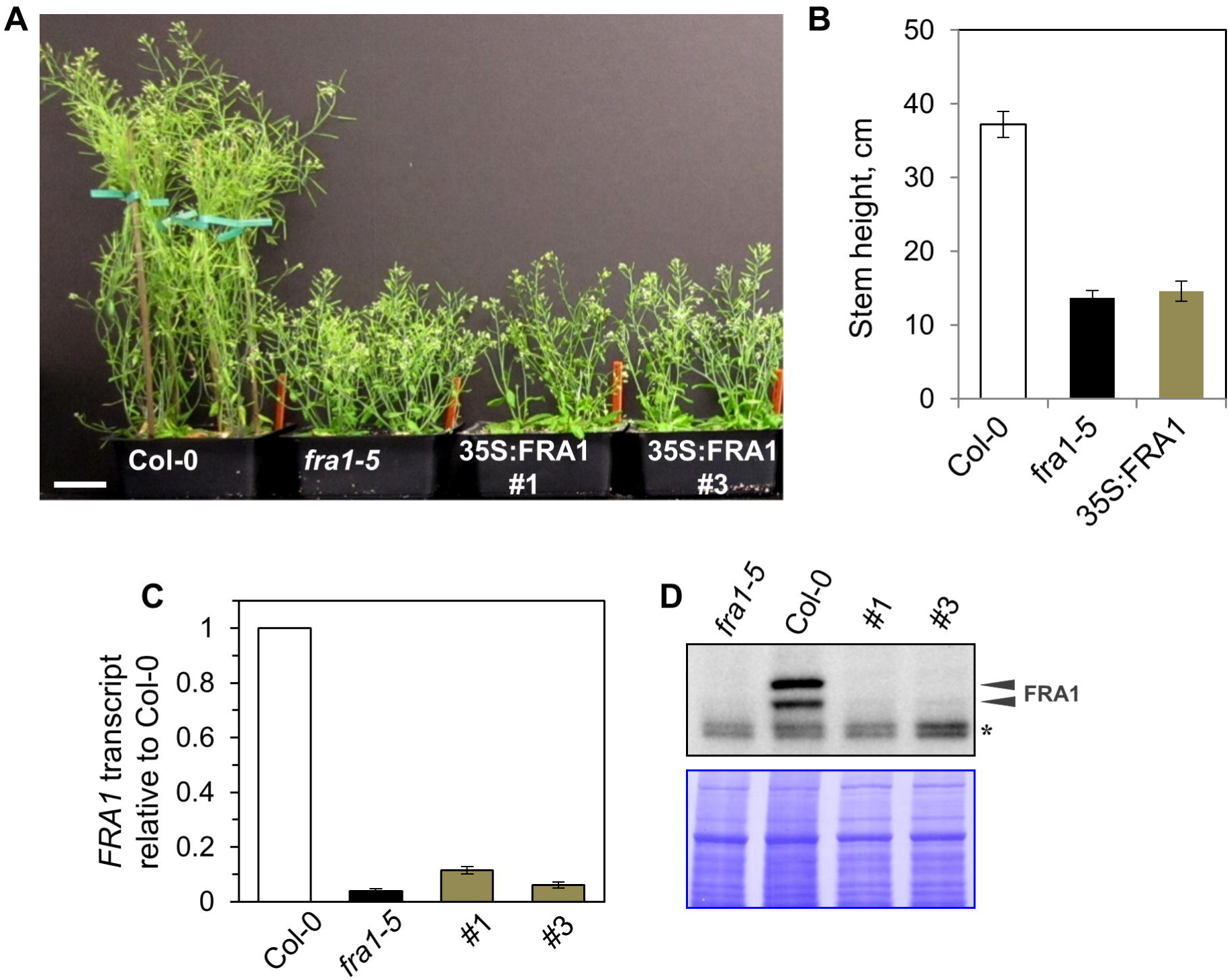

Effect of *FRA1* overexpression in wild-type plants.

(**A**) Images of 4.5-week old plants. Scale bar = 5 cm.

(**B**) Heights of primary inflorescence stems. Values are mean ± SD from at least 8 plants for each genotype.

(**C**) *FRA1* expression level relative to wild-type control. Values are mean ± SEM from three biological replicates.

(**D**) Immunoblot of the native FRA1 protein in *fra1-5*, Col-0 and two independent transformants expressing 35S::FRA1-tdTomato. The native FRA1 protein shows up as two bands (arrowheads). The asterisk marks a non-specific band.

### Summary

Our data show that abundant and processive motility are important for FRA1’s function *in vivo*, consistent with a proposed vesicular trafficking function of FRA1. While several studies have analyzed kinesin processivity mutants *in vitro*, to the best of our knowledge, this is the first study to measure the motility of these mutants *in vivo*. The significant differences between *in vitro* and *in vivo* behavior of the various FRA1 mutants could be due to several possible reasons. Kinesin mutants with positively charged residues introduced in the NL and NCC domains show greater processivity *in vitro* due to electrostatic interactions with the negatively charged carboxy terminus of tubulin under low ionic strength conditions (Shastry and Hancock, 2010; Thorn et al., 2000). In cells, the higher cytoplasmic ionic strength likely disrupts these electrostatic effects thus reducing the gain in processivity. Further, the *in vitro* motility assays are conducted using a truncated kinesin lacking the tail domain and in the absence of cargo and external load. Interactions between the motor and tail domains are critical for regulating kinesin motility (Hammond et al., 2009; Hirokawa et al., 1989), which might account for altered motility of full-length kinesin *in vivo*. Moreover, full-length kinesin transporting cargo in cells will experience load which can promote kinesin stalling and detachment from microtubules (Svoboda and Block, 1994; Valentine and Block, 2009), thus impacting processivity. In addition, regulatory proteins can alter kinesin motility in cells compared to *in vitro* conditions. Therefore, while *in vitro* experiments are helpful to identify potential mechanisms underlying kinesin mechanochemistry, they must be tested *in vivo* to verify their operation under physiological conditions.

## MATERIALS AND METHODS

### Plant materials and growth

Seeds were sterilized with 25% bleach for 10 min, washed with sterile water three times and plated on 0.5X Murashige and Skoog medium (Caisson Laboratories). For live imaging, seeds were stratified for 2-3 days at 4°C and then grown under 16h light for 4 days. For phenotypic analysis, plants were grown in soil under continuous light at 22°C.

### Construct preparation

The FRA1(707)-GFP and proFRA1::FRA1-tdTomato constructs have been previously described in Zhu et al., 2011 and Zhu et al., 2015, respectively. For the 35S::FRA1-tdTomato construct, the FRA1-tdTomato insert was ligated using HindIII and Sal1 restriction sites downstream of the 35S CaMV promoter in pCAMBIA1300. For the processivity mutants, site-directed mutagenesis was conducted using the megaprimers listed in Supplemental Table 1.

### Quantitative RT-PCR and immunoblotting

RNA and proteins were isolated from 2-week old seedlings. Total RNA was extracted by the Trizol method and complementary DNA (cDNA) synthesized with revertAid First Strand cDNA Synthesis Kit (ThermoFisher Scientific). Quantitative RT-PCR was performed using the SYBR method with primers listed in Zhu et al., 2015.

For immunoblotting, lysates were prepared by grinding seedlings in liquid N_2_ and homogenizing in protein isolation buffer (50 mM Tris-acetate, pH 7.5, 2 mM EDTA, 1 mM PMSF and protease inhibitor tablet from Roche). Total proteins (~30 μg each) were separated by SDS-PAGE, transferred to PVDF membrane and probed with anti-FRA1 primary antibody (1:1,000, Zhu et al., 2015) and anti-rabbit HRP secondary antibody (1:5,000, Jackson Immuno Research).

### Protein purification and *in vitro* motility assays

His-tagged FRA1(707)–GFP was expressed in BL21(DE3) Rosetta *Escherichia coli* using 0.5 mM IPTG at 24°C for 4 h and purified using Ni–NTA resin affinity chromatography (Qiagen). The purified fusion protein was then desalted using a PD-10 column (Amersham Biosciences) and exchanged into MAB buffer (10 mM PIPES, 50 mM potassium acetate, 4 mM MgSO4, 1 mM EGTA, pH 7.0) supplemented with 50 mM NaCl, which has an ionic strength of about 110 mM. Single-molecule motility assays were carried out exactly as described in Zhu et al., 2011.

### Live imaging and data analysis

Live imaging of FRA1-tdTomato was carried out in apical hypocotyl cells of four-day old seedlings using variable angle epifluorescence microscopy. Specimens were excited with 3 mW 561-nm laser (Melles Griot) and images gathered using a 100x (N.A. 1.45) objective at 1-s intervals with an EM-CCD camera (ImagEM, Hamamatsu). Kymograph analysis was used to measure the speed and run length for individual motile events. Mean speed was determined by Gaussian fitting using the least-squares method to the measured speed distribution. For mean run length, cumulative distribution functions (CDFs) were calculated from the distributions of measured run lengths and a monoexponential function was fitted to 1-CDF data using the least-squares method.

## COMPETING INTERESTS

The authors declare no competing and financial interests.

## AUTHOR CONTRIBUTIONS

R.D. and A.G designed the study and experiments. A.G generated the constructs and did the *in vitro* and *in vivo* experiments. L.D performed the qRT-PCR experiments. A.G and R.D analyzed the data and wrote the manuscript.

## FUNDING

This work was supported by NSF grant MCB-1121287 to R.D. and by the NSF Science and Technology Center for Engineering Mechanobiology, award number CMMI-1548571.

